# Major changes in gene expression between strains of diapausing and non-diapause spruce budworm despite little evidence of genetic divergence

**DOI:** 10.1101/2024.11.20.624545

**Authors:** Karin R. L. van der Burg, Jeff Gauthier, Amanda D. Roe, Michel Cusson, Brian Boyle, Roger C. Levesque, Katie E. Marshall

## Abstract

To survive harsh winter conditions, many insects enter a state of dormancy called diapause which allows them to withstand extreme temperatures and lack of food. When diapause is induced, insects enter a state of arrested development and low metabolic rate. Because diapause and environmental conditions are closely linked, variation within and between species in diapause induction, depth, and duration is extremely common. Studies investigating the genetic underpinnings of diapause tend to focus on either different populations and/or environmental variation, which runs the risk of confounding the genetic signal either due to isolation-by-distance or variation in environmental conditions. We use the eastern spruce budworm (*Choristoneura fumiferana*) to circumvent these issues and investigate within-population variation in diapause. *C. fumiferana* usually diapauses as second instar larvae, but a small subset of individuals does not diapause and instead continues through development. This non-diapause phenotype can be selected to create a non-diapause strain. Here, we present a chromosome-level assembly of such a non-diapause strain, and compare it to an earlier published genome assembly of the diapause strain. We did not find evidence of major chromosome rearrangements, indicating that the genetic variation between strains is likely small and multi-genic. Gene expression comparisons between the strains indicate major gene expression changes, where genes associated with glycolysis and environmental signaling processing increase in expression in the diapause strain. Lastly, we found that gene expression diverges halfway through the first instar, and not before, indicating that the signal to induce diapause happens early in the first instar.

**Significance statement:** The ability to enter diapause is a crucial adaptation for many insects to survive cold winters. The genetic underpinnings of the diapause phenotype has been the target of many investigations. However, many of these studies focus on variation in diapause on between geographic regions or changes in environmental conditions, which can lead to confounding factors. Here, we focus on within-population variation in diapause in the eastern spruce budworm, with two strains established from the same source population that vary in their ability to induce diapause. We present a chromosome level assembly of the non-diapause strain, and report that genetic changes between the diapause and the non-diapause line are likely small and involve multiple loci. Gene expression differences were extensive, and mostly associated with mechanisms to avoid freezing. This work sets the stage to further investigate how minimal genetic changes can lead to large-scale phenotypic changes as impactful as diapause.

## Introduction

Insects living in temperate climates often face stressful seasonal conditions where extreme temperatures and lack of food are common. One adaptation to winter conditions is diapause, a hormone-induced dormant state associated with extensive physiological changes such as arrested development and low metabolic activity (Gill et al., 2017). When diapause is induced, insects shift away from a direct developmental program and start to prepare for diapause through changes in gene expression, neuroendocrine signaling, and metabolism until diapause is initiated (Denlinger, 2002; Kostál, 2006). Most diapausing insects have a responsive diapause program, where an external signal such as photoperiod or temperature during a sensitive period is translated to an internal signal to induce the diapause program (Wilsterman et al., 2021), although species with an obligate diapause, which is induced regardless of external conditions, do exist (Kostál, 2006). Because diapause and environmental conditions are so closely linked, there is often extensive local adaptation in diapause regulation (Pruisscher et al., 2018; Tyukmaeva et al., 2011). Thus, studies investigating genetic mechanisms underlying diapause tend to focus on either between-population variation (Fabian et al., 2012; Kozak et al., 2019), where diapause is locally adapted, or within-population comparisons of responsive diapause programs, where environmental conditions vary to induce or prevent diapause (Ragland et al., 2010). While useful and informative, studying between-population variation risks confounding genetic signals due to isolation by distance with natural selection on population-specific traits. For within-population comparisons of responsive programs, it can be difficult to disentangle the diapause-specific mechanisms from general environmental variation in the response mechanisms. In contrast, within-population genetic variation in obligate diapause induction should allow for the investigation of the genetic changes necessary to alter the diapause program without confounding effects from isolation-by-distance or variation in environmental conditions.

Diapause is a complex trait, and significant research has been conducted to resolve its underlying regulation and genetic architecture (Denlinger, 2022; Ragland et al., 2019). Given that diapause has evolved multiple times, it stands to reason that the genetic basis of this trait will be equally varied. Several studies have implicated circadian clock genes in diapause variation along a latitudinal gradient, such as in the European corn borer *Ostrinia nubilalis* (Kozak et al., 2019), the parasitic wasp *Nasonia vitripennis* (Dalla Benetta et al., 2019), the speckled wood butterfly *Pararge aegeria,* and several other species (Mathias et al., 2007; Tauber et al., 2007). The identification of circadian clock genes associated with clinal changes in diapause is unsurprising, as photoperiodism is the most reliable environmental cue to predict seasonal changes, and also varies significantly with latitude. A few classic studies in *Drosophila melanogaster* identified different signaling genes underpinning diapause regulation: insulin-regulated *Dp110* (Williams et al., 2006), as well as *couch potato*, a gene believed to be involved with ecdysteroid signaling (Cogni et al., 2014). Although the link with environmental conditions is less clear-cut, both insulin and ecdysteroids are known to be environmentally responsive endocrine signals. However, all of these studies were conducted on insects from different populations, where isolation-by-distance and/or environmental variation play a huge role in the genetic divergence between populations.

The eastern spruce budworm, *Choristoneura fumiferana* (Clemens, 1865, Lepidoptera:Tortricidae), is a widespread conifer pest that enters what is usually considered an obligatory diapause to survive winter throughout its range. Diapause is induced in early instar larvae during late summer and early fall in anticipation of stressful winter conditions, regardless of external conditions. The process of diapause induction and preparation is complex and temperature-sensitive (Han and Bauce, 1998; Marshall and Roe, 2021; Roe et al., 2024). Typically, *C. fumiferana* is described as univoltine, with one single obligatory diapause (Marshall and Roe, 2021). After hatching, larvae forego feeding, and seek an overwintering site on the host plant where they construct a silken structure (hibernaculum) prior to molting and spinning a second hibernaculum layer before becoming dormant (Régnière and Duval, 1998). However, variation in diapause expression exists within this species (Fig. 1). At one extreme, additional diapause stages may occur following emergence in spring, requiring two years for an individual to complete development (Harvey, 1961). At the other end of this continuum, diapause free development (non-diapause) was documented in multiple laboratory rearings of *C. fumiferana* (Harvey, 1957). Here, second instar larvae were able to continue development and did not require a period of developmental arrest. Interest in this non-diapause phenotype led Harvey to develop a non-diapause colony of *C. fumiferana*. He used continuous light to select for non-diapause development from *C. fumiferana* families that showed elevated expression of this phenotype. After 12 generations of selection, this colony showed nearly complete non-diapause expression. Because the selection experiment showed such rapid change, it can be assumed that artificial selection acted on standing genetic variation within this population, rather than novel mutations (Harvey, 1957). Furthermore, artificial selection is the strongest possible form of natural selection for one specific phenotype (non-diapause), and so we expect the genetic loci selected for in the non-diapause strain to all be associated with the non-diapause phenotype. Thus, *C. fumiferana* is an ideal model system for examining the genetic architecture and regulatory mechanisms of diapause without confounding effects of population structure or environmental differences.

**Figure 1:**
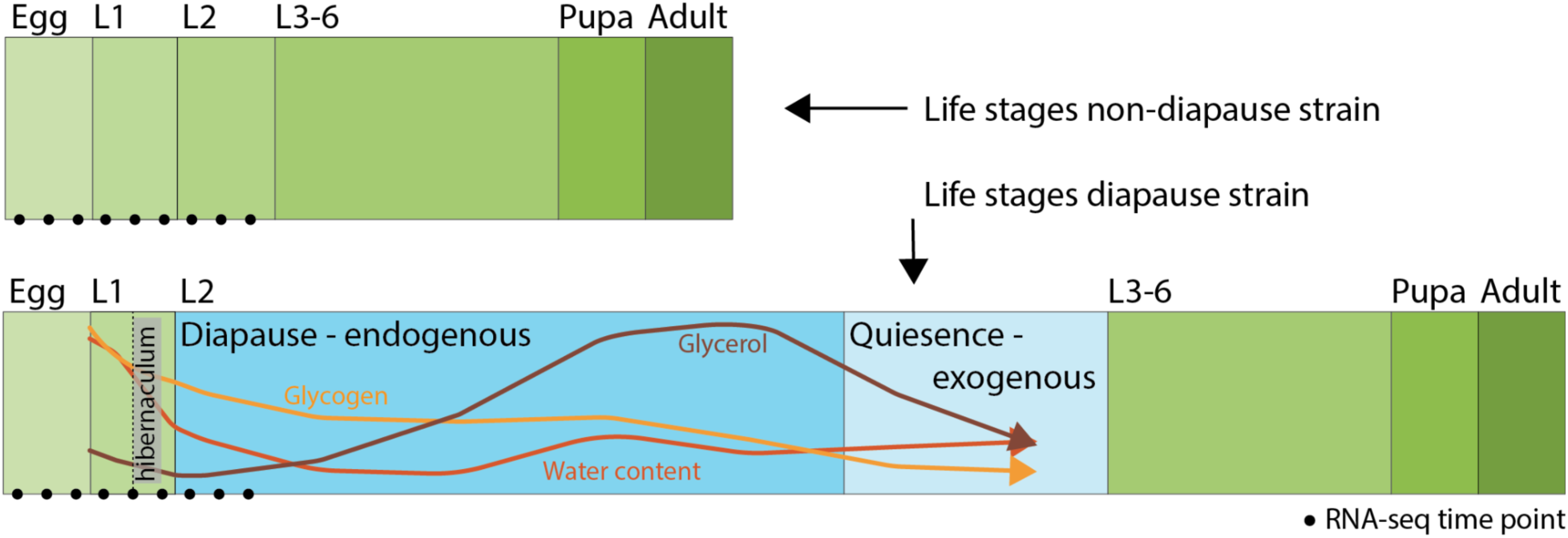
Life stages for the two strains of spruce budworm. Spruce budworm experience periods of active development (green) and developmental arrest (blue). The period of developmental arrest in the diapause strain is an obligatory diapause that occurs during the 2nd larval instar. During diapause changes in glycerol, glycogen and water content are shown (Han and Bauce, 1998).

The availability of a non-diapause colony and genomic tools provides an ideal opportunity to examine the genetic basis of diapause in *C. fumiferana*. Recently, a chromosome-level assembly of the diapause strain of the spruce budworm was published (Béliveau et al., 2022). Here, we present a chromosome-level genome assembly of a non-diapause strain of *C. fumiferana*, as well as a gene expression comparison with fine temporal resolution between the diapause and the non-diapause strain. Although we do not see evidence of major chromosomal inversions or significant SNP pileups, we did find extensive changes in gene expression between the diapause and non-diapause strain starting midway through the first larval instar. This work sets the stage for future investigation into the *C. fumiferana* diapause phenotype.

## Materials and Methods

### Study organisms and tissue sampling

#### Non-diapause laboratory colonies

Non-diapause *C. fumiferana* were sampled from two strains for this work. The original Harvey (1957) strain was reared continuously for 60 years at the Insect Production and Quarantine Laboratories following standard rearing protocols (Ebling and Dedes, 2015) with the modification of egg masses being put directly on to the McMorran diet for continuous development (McMorran, 1965). This original strain, along with a wild-type (=diapausing) laboratory colony (Glfc:IPQL:Cfum; Roe et al. 2018), were used for the comparative RNA-seq analyses described below. A second non-diapause strain was again developed from the diapausing laboratory strain using similar selection protocols (Harvey, 1957) after the loss of the original non-diapause strain. Individuals from this second strain were collected for whole genome sequencing, Omni-C, and long read transcriptomics.

#### DNA extraction for genome sequencing

High molecular weight DNA was extracted from a 3–4-day old male *C. fumiferana* pupae obtained from the Insect Production and Quarantine Laboratories (Great Lakes Forestry Centre, Sault Ste. Marie, ON, Canada). Pupae were dissected on ice to remove the gut tissue prior to processing with the MagAttract HMW DNA kit (Qiagen, 67563) following the manufacturer’s “Fresh or Frozen Tissue” protocol, where we replaced all vortexing steps with gentle inversions to minimize DNA fragmentation. DNA was quantified with the Qubit dsDNA BR assay on a Qubit 2 fluorometer (Thermo Fisher), then evaluated for purity with a NanoDrop 2000 spectrophotometer (Thermo Fisher) with A260/A280 and A260/A230 ratios above 1.8 as thresholds.

#### Omni-C library preparation

A single 3-4 day old pupae was used for Omni-C library preparation using the Dovetail (R) Omni-C ® kit (Cantata Bio, Scotts Valley, CA, USA) following manufacturer’s instructions (Omni-C protocol, non-mammalian samples v1.0, Insects & marine invertebrates). The library was sequenced at the Centre d’Expertise et de Services Génome Québec (Montreal, QC, Canada) on an Illumina Novaseq shared S4 lane.

### Transcriptomics

#### Short-read transcriptomics

For gene annotation and gene expression comparison between the two diapause phenotypes, we used comparative transcriptomic data from Béliveau (et al., 2022) that sampled early life stages (i.e., eggs, 1st, and 2nd instar larvae) from a diapausing *C. fumiferana* strain (Roe et al., 2018) and the original non-diapause strain (Harvey 1957). In short, both colonies were reared under standard conditions (16h:8h Light:Dark; 23°C; 60-70% relative humidity) using standard protocols (Ebling and Dedes, 2015). They extracted RNA from a pool of 10-60 eggs per time point (sampled 1, 3 and 5 days post-oviposition), a pool of 10-60 1st instar larvae per time point (sampled 1, 3, and 5 days post-hatch) and a pool of 10-60 2nd instar larvae per time point (sampled 1, 3, and 5 days post-molt). Later life stages were also sampled from both strains, including 6th instar larval heads (day 3 post-molt, pool of 20 individuals), two male and two female 5th instars immediately prior to molting, and two males and two female 6th instars (sampled day 2 post-molt). From adult moths, data was reported for male and female antennae, brain, and reproductive tract, as well as the female pheromone gland.

#### Long read transcriptomics

The whole transcriptome of a 6th instar non-diapause male larva was extracted with the TRIzol method (Rio et al., 2010) using Thermo Fisher’s TRIzol Reagent. Cell tissues were flash-frozen in liquid nitrogen and ground using a mortar and pestle; afterwards, the manufacturer’s recommended protocol was used. Total RNA was quantified with the Qubit RNA HS kit (Thermo Fisher), and purity was assessed with a NanoDrop 2000 spectrophotometer (Thermo Fisher). RNA was considered pure if A60/A280 and A260/A230 absorbance ratios were above 1.8. Total RNA was then sequenced natively (i.e. without the need for reverse transcription of ribosomal RNA depletion) with Oxford’s Nanopore’s direct RNA sequencing procedure (SQK-RNA004) on a R9.4.1 flow cell (FLO-MIN106). Indeed, direct RNA sequencing adapters bind to poly-A tails in 3’, thereby selectively binding to eukaryotic messenger RNA (SQK-RNA004).

### Whole Genome Sequencing

Three Oxford Nanopore Technologies (ONT) MinION sequencing runs were pooled to 18.6 gigabases, about 25X coverage assuming a genome size of 600 Mb (Béliveau et al., 2022). For each run, 1 µg of pure genomic DNA was processed with the Ligation Sequencing Kit (SQK-LSK109 or SQK-LSK110). The first run was done on a R9.4.1 flow cell (FLO-MIN106). The second run, also on a R9.4.1 flow cell, was run as previously described, but with DNA treated by the Short Read Eliminator XS size-selection kit (Pacific Biosciences). The third run was as the second one but with a R10.4.1 flow cell (FLO-MIN110). These three sequencing runs yielded 6.5 Gb, 8.5 Gb and 3.1 Gb respectively. In addition, a whole genome shotgun library was generated with ∼500 ng total genomic DNA using the NEBNext Ultra II kit (New England Biolabs). Prior to library preparation, the DNA was sheared to an average size of 600 bp using a Covaris M220 (Covaris). Sequencing was done on a MiSeq apparatus using a 600 cycle v3 kit (2 x 300) at the Plateforme d’Analyses Génomique (Institut de Biologie Intégrative et des Systèmes, Université Laval, Quebec, Canada), leading to 31.6 million read pairs (19.1 Gb in total).

### Draft hybrid assembly

Both long and short read sets were assembled in a long-read-first hybrid assembly approach. Briefly, ONT reads were quality-filtered with NanoFilt v2.6.0 (De Coster et al., 2018) to select only reads above a mean Phred score of 10 and a minimum length of 4000 bp. Likewise, Illumina MiSeq reads were processed with TRIMMOMATIC v0.39 (Bolger et al., 2014) to only select reads above Q20 over at least 36 bp. Filtered ONT reads were assembled de novo with CANU v1.9 (Koren et al., 2017) using default parameters and an estimated genome size of 600 Mb. The final contigs draft from CANU was polished by aligning the Illumina MiSeq reads using Pilon v1.23 (Walker et al., 2014). This step was repeated three times, i.e. until no more draft bases were corrected by Pilon. A final scaffolding step was run with LongStitch v1.0.2 (Coombe et al., 2021) to cut and re-scaffold the draft assembly at potentially misassembled regions.

### Omni-C scaffolding

An in-house pipeline was developed to pre-process Omni-C reads, remove internal adapters, and identify true Omni-C contacts usable for bridging contigs. First, Omni-C reads were trimmed with TRIMMOMATIC v0.39 (Bolger et al., 2014) to select only reads above Q30 over a minimum 50 bp length. Then, CutAdapt v3.2 (Martin, 2011) was used to remove the internal adapter bridging the crosslinked Omni-C fragments. The adapter sequence itself was not provided publicly by Dovetail Genomics but was nevertheless identified by analyzing the k-mer frequency among a subset of merged Omni-C read pairs. A 30-mer internal palindrome was found in the top three 31-mers of this subset.

The cleaned Omni-C reads were then aligned to the hybrid draft assembly with Bowtie2 using the preset (--very-sensitive-local) to prevent the loss of Omni-C mappings due to small mismatches and gaps). The resulting BAM files were processed with HiCExplorer v3 (Wolff et al., 2020) to build a Hi-C contact matrix devoid of problematic read pairs (one mate not unique, one mate low quality, one mate unmapped, duplicated pairs, self-ligation, etc.). Given that Omni-C uses a non-specific DNase instead of a restriction endonuclease (“Omni-C 0.1 documentation,” n.d.), the cut sites and dangling sites were set to “NNNN”. Reads used in the final contact matrix were output to BAM format, then used for scaffolding the Canu-Pilon-LongStitch draft with SALSA2 (Ghurye et al., 2019), with the genome size (--exp) set at 580 Mb and the cutting enzyme (--enzyme) set at “DNASE”. This procedure was repeated three times, for which contigs below 50 kb, 100 kb and 120 kb were filtered out of the draft at each iteration.

### Final scaffolding and gap filling steps

The Omni-C scaffolded draft was further scaffolded by use of the wild-type *C. fumiferana* genome sequence (Béliveau et al., 2022) with RagTag version 2.1.0 (Alonge et al., 2022). Briefly, RagTag corrects probable misassemblies by comparing the draft to a reference, scaffolds the draft over the reference, uses Hi-C information to resolve conflicts and patches gaps. After this step, a final Omni-C scaffolding step occurred as described in the paragraph above, this time without a contig length cutoff applied to the input draft.

### Genome assembly and annotation

Genome completeness was verified using BUSCO (Manni et al., 2021). To annotate the genome, we used the MAKER annotation software (Holt and Yandell, 2011). In the first round, we used a spruce budworm transcriptome, repeat library and known proteins to inform gene predictions. To create a repeat library, we used repeatmodeler to identify repeats in the genome and create a repeat library. For the transcriptome, we used the same RNA-seq data as reported in Béliveau et al. (2022). Long RNA-seq reads were mapped using minimap2 (settings: -ax -k14 splice). All RNA data was assembled using Trinity using the --long_reads_bam setting to account for long reads in the data (Grabherr et al., 2011). Lastly, we used the protein library from the diapause strain of the spruce budworm as additional input for the first rounds of gene predictions. In the second and third rounds of MAKER iteration, we used both augustus and SNAP training software to improve gene annotations (Korf, 2004; Stanke and Morgenstern, 2005). For functional annotations of our gene model we used interproscan to scan for homology against the uniprot protein database (Blum et al., 2021; Jones et al., 2014; The UniProt Consortium et al., 2023).

### Genome comparison

To test whether major chromosome rearrangements had taken place between the diapause strain (Genbank reference: GCA_025370935.1 (Béliveau et al., 2022)) and the non-diapausing strain, we used *Satsuma* (Grabherr et al., 2010) to align the respective genomes. The resulting alignments were visually inspected for major rearrangements. The same analysis was done using *mummer* (Kurtz et al., 2004).

In addition, we called nucleotide variants between the two genomes and looked for elevated SNP occurrences in a sliding window analysis (20k wide, with 10k sliding steps), where the number of variants were counted in each window. We then ranked windows according to variant frequency, and widows with a high frequency of variants that did not appear to be associated with repetitive elements were inspected further.

### Differential gene expression

To compare gene expression between the diapausing and non-diapause strains, we used the mRNA-seq data from eggs, L1 and L2 instar larvae as described previously. We investigated which genes were consistently differentially expressed between the diapausing and the non-diapause strain. mRNA reads were aligned to the non-diapause genome, and reads within gene annotations were counted using *featureCounts* from the R package Rsubread (Liao et al., 2019). We then used the software *Short Time-series Expression Minor* (*STEM*, (Ernst and Bar-Joseph, 2006)) to analyze gene expression. We used *STEM* to cluster gene expression patterns in 20 different temporal expression profiles, following a log2 normalization. To calculate which profiles have significantly more genes assigned then expected, a permutation test was performed: original time point values are randomly permuted, expression values are renormalized, and genes were assigned to most closely matching profiles, and repeated 50 times, to determine the expected number of genes for each profile. Significance was determined after a Bonferroni correction for multiple hypothesis testing, and was based on the total number of genes assigned and the expected number of genes assigned. We then determined GO-term enrichment for each of the significant profiles. We used *Revigo* (Supek et al., 2011) to cluster and summarize GO terms to find the representative GO terms.

Next, we compared temporal expression profiles across the different strains. For each non-diapause profile, we determined the number of genes assigned to a different profile in the diapause strain–the intersection of two profiles. Significance of the number of genes in the intersection was computed using the hypergeometric distribution based on the number of total genes assigned to each of the two profiles, and the total number of genes in the experiment. Thus, we identified which gene groups showed different temporal expression patterns, and which GO-terms were enriched in those groups. We used *Revigo* (Supek et al., 2011) to cluster and summarize GO terms to find the representative GO terms.

## Results

### Genome assembly and annotation

The genome assembly resulted in 30 full-length scaffolded sequences larger than 6 Mbp, representing a chromosome level assembly. 769 smaller scaffolds were also assembled, with a size ranging from 200 base pairs to ∼450 Kbp. Busco analysis indicated a 98.5% completeness score (complete + fragmented), of which 3.4% was duplicated. 42.6% of the genome was soft-masked due to the presence of repetitive sequence. The MAKER annotation pipeline of the non-diapause genome resulted in 21,609 gene annotations with 91% of the genes showing an AED-score of 0.5 or better.

**Table 1.** Comparison statistics for the diapause and the non-diapause genome.

|  | *diapause* | *non-diapause* |
| --- | --- | --- |
| *N50* | *20.6 Mbp* | *18.5 Mbp* |
| *Assembly size* | *575.3 Mbp* | *572.3 Mbp* |
| *Scaffolds (>6 Mbp)* | *30* | *30* |
| *Mbp captured in scaffolds > 6 Mbp* | *569.6 Mbp* | *521.4 Mbp* |
| *Scaffolds* | *334* | *799* |
| *Mbp captured in scaffolds < 6 Mbp* | *5.7 Mbp* | *50.9 Mbp* |
| *Busco score (complete + fragmented)* | *97.7%* | *98.50%* |
| *Busco score duplicated* | *3.9%* | *3.40%* |

### Genome comparisons of the diapause and non-diapause strain

Genome alignment did not show any evidence of major chromosome rearrangements between the diapausing and non-diapausing strain (Figs. 2, S1). We then compared nucleotide variation between the genomes of the two strains to identify concentrated regions of genetic divergence between the strains. Although elevated levels of SNP variation were found on chromosomes 10 and 30, these regions also contained elevated levels of repetitive sequences or had missing nucleotides (Ns) nearby. Both features could artificially increase the number of SNPs due to assembly artifacts.

**Figure 2:**
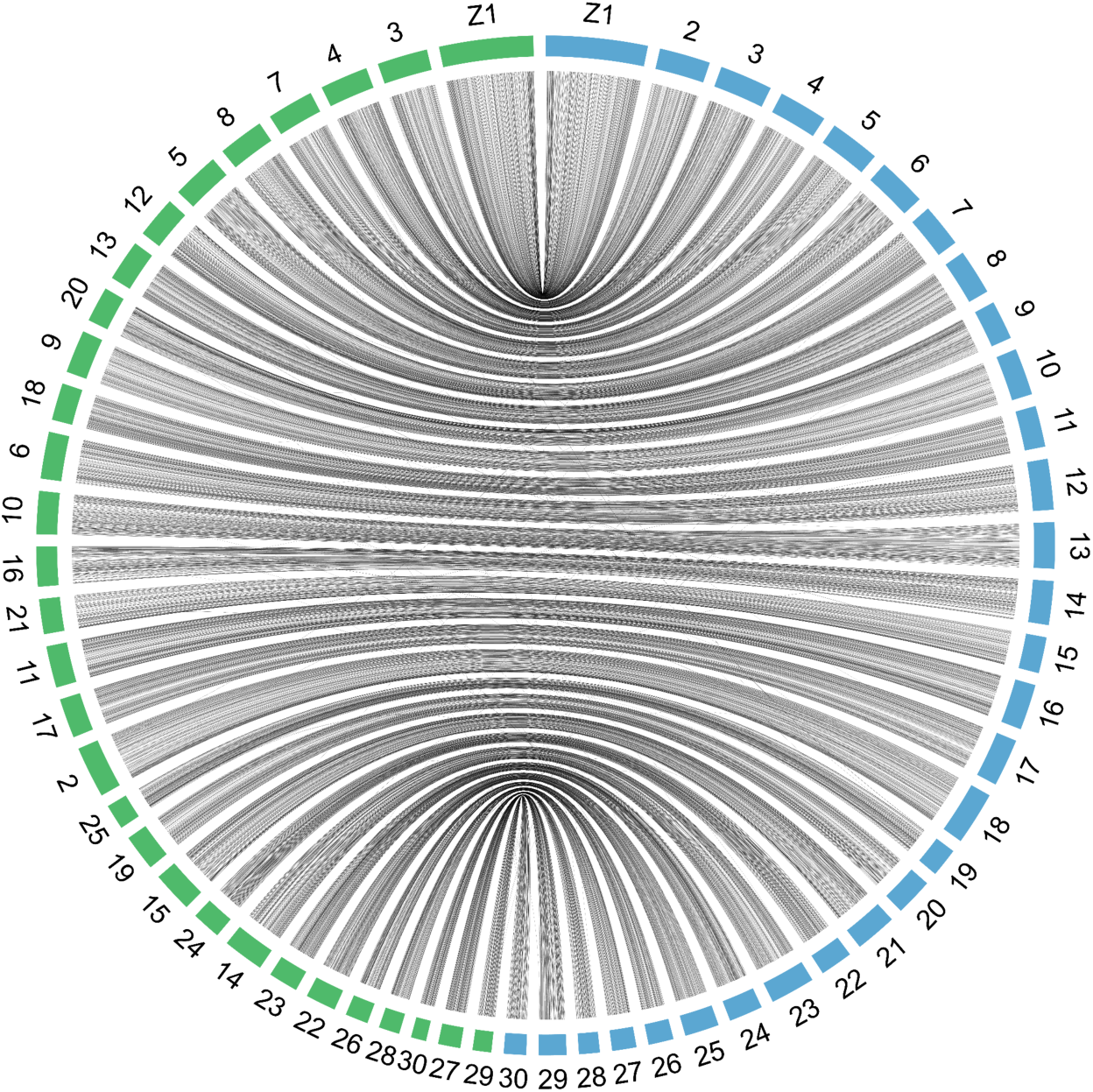
Chromosomal alignment between the diapausing (blue) and non-diapause strain (green). Alignment of two genomes reveals high levels of similarity between genomes. There is no evidence of major chromosomal rearrangements between the two strains.

### Differential gene expression

We analyzed temporal gene expression profiles between the early life stages of the diapause and non-diapause strains, which corresponds to the time when diapause is induced. First, we identified significantly enriched profiles for each of the two strains separately. We assigned expression profiles for each gene to 20 different clusters and found that both the diapause and the non-diapause strains had five temporal expression profiles (Fig. 3) with significantly more genes assigned than expected per a permutation test (Table 2). Go-term enrichment was determined by dividing the number of genes assigned by the number of genes expected. The highest fold-enriched GO-term was displayed in Fig 3. For a complete list of GO-terms enriched in each profile, see table S1.

**Figure 3:**
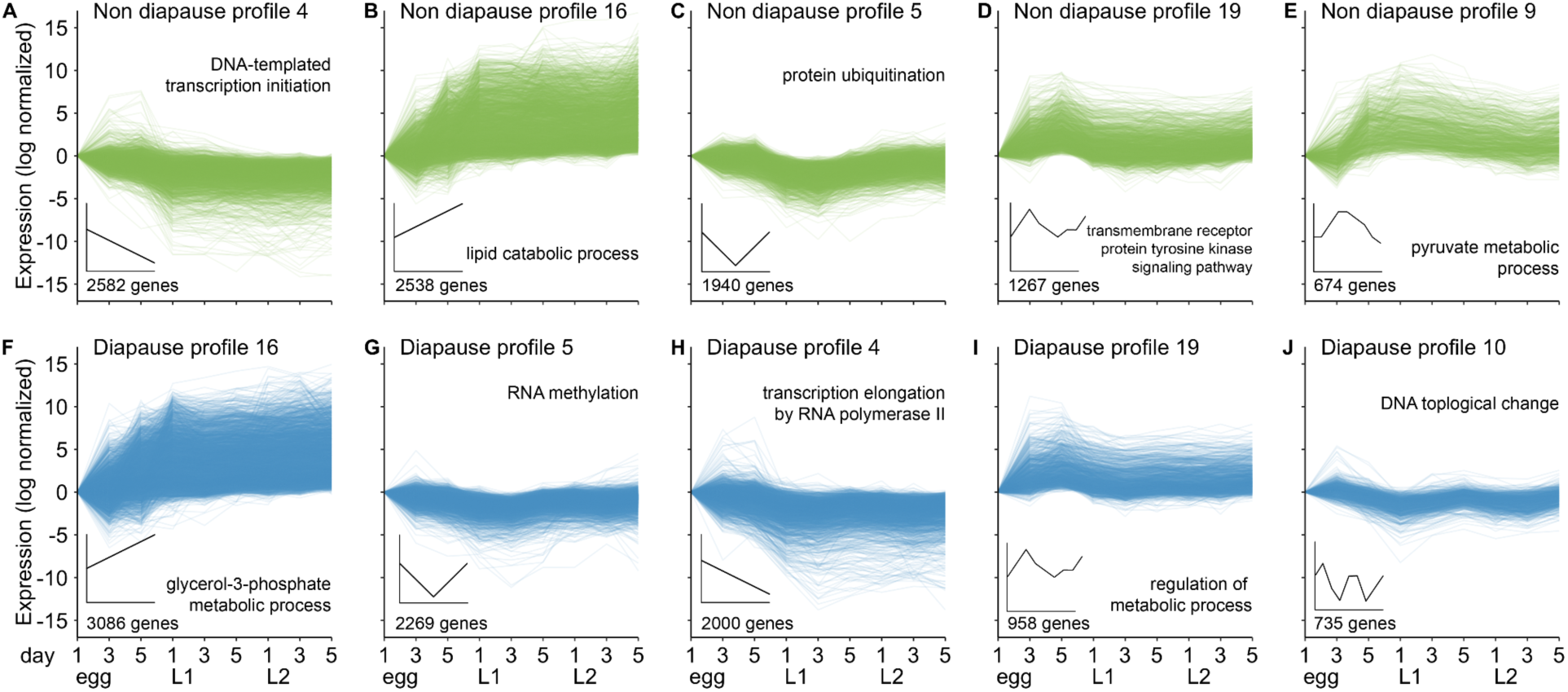
Temporal gene expression profiles for the diapause and non-diapause strain Spruce budworm larvae were sampled over nine days, from three different life stages: egg, L1 and L2 for each strain; non-diapause (A-E) and diapause (F-J). Similar temporal gene expression profiles were clustered together (black inserts display general shape of expression profile). For each strain, five expression profiles had significantly more genes assigned than expected based on a permutation test (displayed here). The number of genes and the top enriched GO-term for each profile are shown.

**Table 2.**
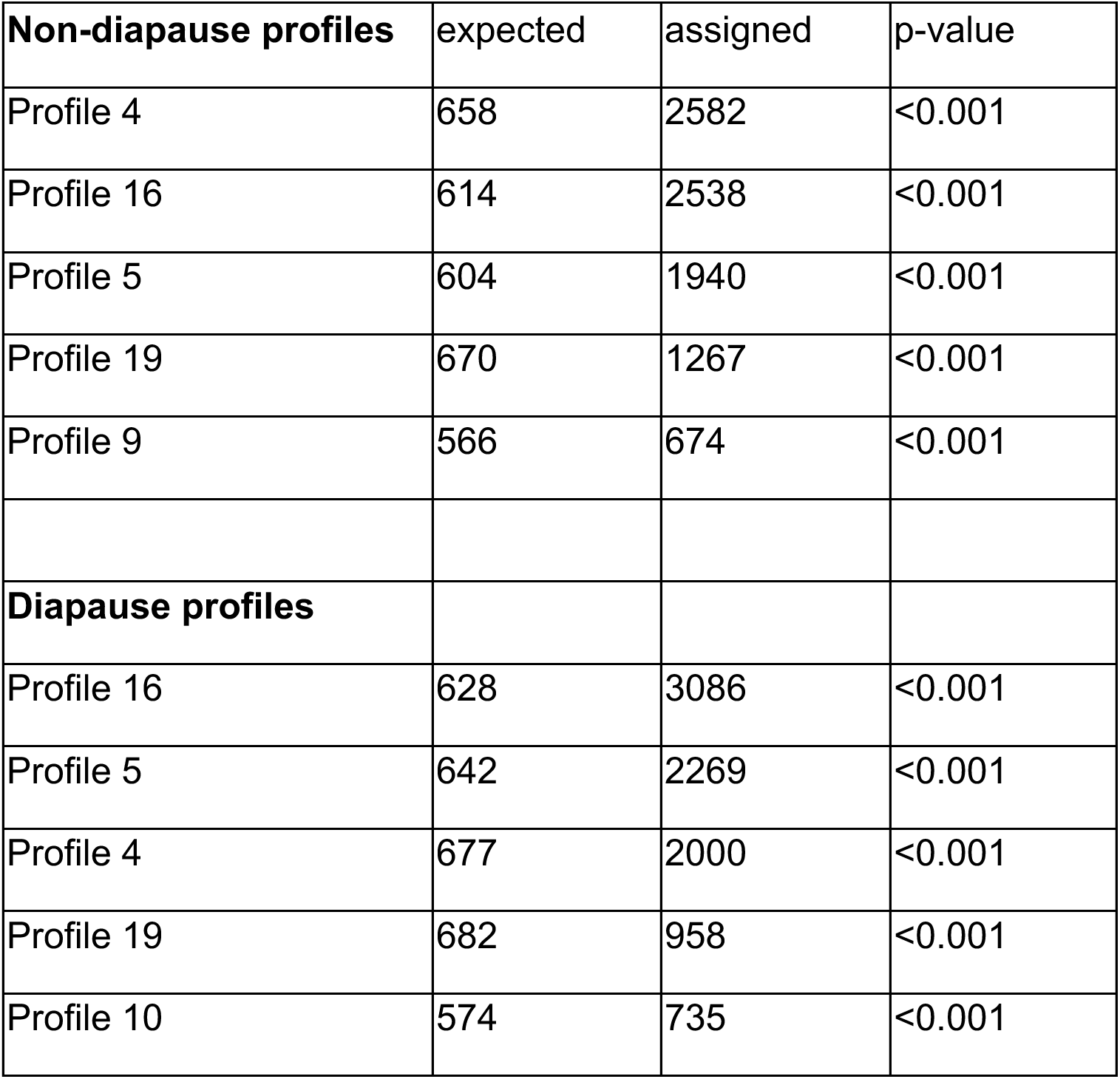
Expected number of genes versus assigned number of genes. Expected number of genes is based on a permutation test.

| **Non-diapause profiles** | expected | assigned | p-value |
| --- | --- | --- | --- |
| Profile 4 | 658 | 2582 | <0.001 |
| Profile 16 | 614 | 2538 | <0.001 |
| Profile 5 | 604 | 1940 | <0.001 |
| Profile 19 | 670 | 1267 | <0.001 |
| Profile 9 | 566 | 674 | <0.001 |
| **Diapause profiles** |  |  |  |
| Profile 16 | 628 | 3086 | <0.001 |
| Profile 5 | 642 | 2269 | <0.001 |
| Profile 4 | 677 | 2000 | <0.001 |
| Profile 19 | 682 | 958 | <0.001 |
| Profile 10 | 574 | 735 | <0.001 |

We then used the expression profiles of the non-diapause strain as a baseline to determine which genes in the diapause strain showed divergent expression patterns. Although our experimental design does not allow for direct differential gene expression comparison due to lack of time point replication, we can identify contrasting expression profiles over time and identify key gene expression differences between the two strains. For three of the five gene expression profiles in the non-diapause strain (Profiles 4, 5, and 9), genes in the diapause strain showed significantly different expression profiles, with more than 10% of divergent genes (Figures 4-6). We considered these differences to represent genes that could be associated with regulating the diapause phenotype.

**Figure 4:**
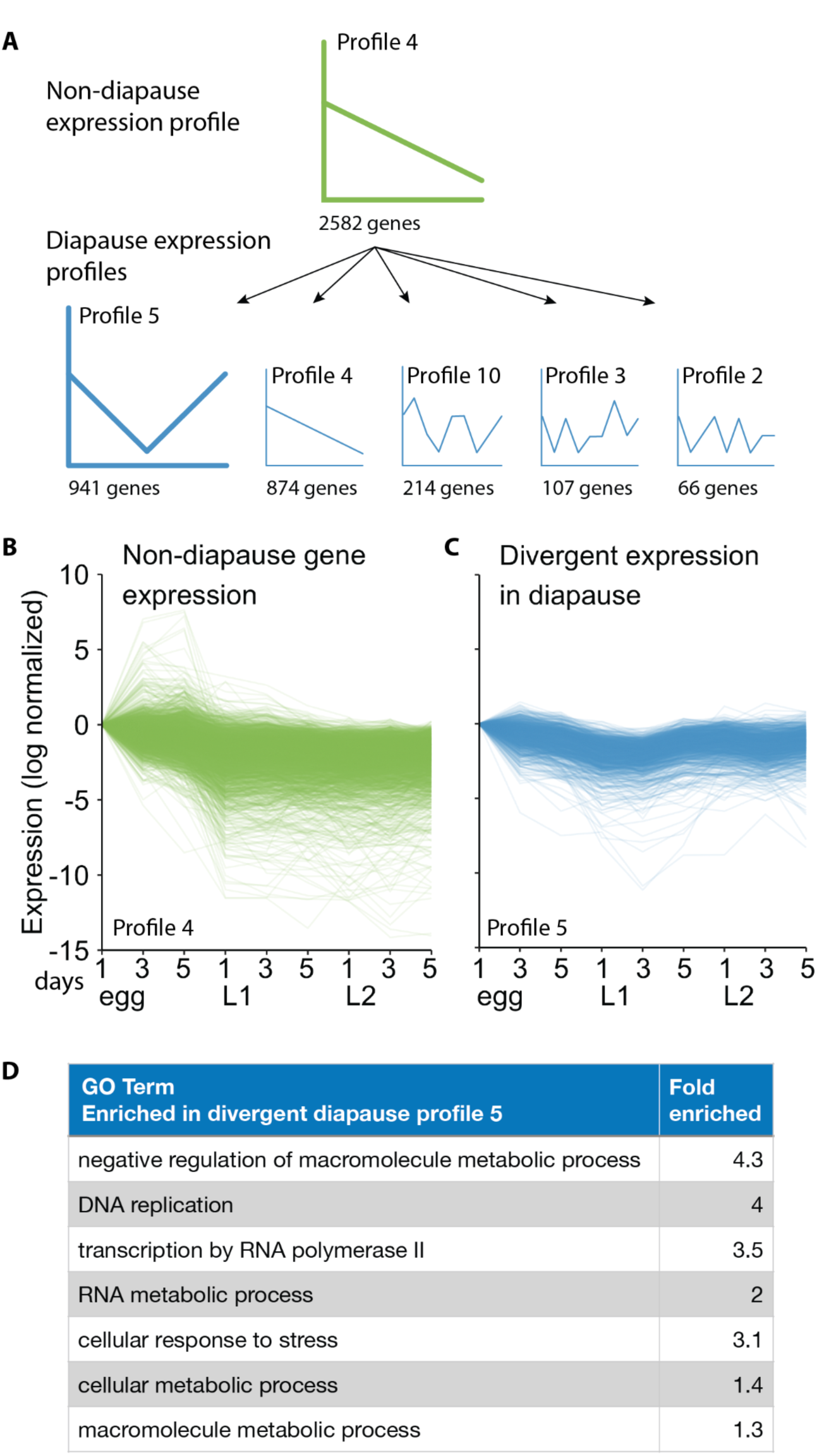
Divergence between the non-diapause temporal gene expression profile 4 and the diapause strain. A. Genes assigned to temporal expression profile 4 in the non-diapause strain were assigned to different profiles in the diapause strain. B. Non-diapause profile 4. C. Divergent genes assigned to diapause expression profile 5. D: GO term enrichment in the divergent gene expression in profile 5. Fold enriched is the total number of genes over the expected number of genes assigned to a GO term.

In non-diapause profile 4, 2582 genes had expression that continually decreased throughout the sampled timepoints (Fig. 4A-B). When contrasted with the non-diapause strain, 874 genes in the diapause strain showed the same expression profile, while 941 genes increased in expression midway through the 1st instar (i.e. Profile 5) (Fig. 4C). Diapause profiles 10, 3 and 2 had 214, 107, and 66 genes assigned to them, respectively (Fig. 4A). GO-term enrichment in the largest differing profile was associated with the negative regulation of developmental processes and DNA replication (Fig. 4D for a complete list). Genes assigned to each GO-term are reported in Table S2.

In non-diapause Profile 5, 1940 genes exhibited expression that decreased during egg and early 1st instar development, but increased in expression by the 2nd instar (Fig. 5A,B). Of those genes, 482 genes were assigned to the same temporal expression profile (i.e. Profile 5) in the diapausing strain. However, 824 genes showed a different temporal expression profile, where expression continued to decline through the 2nd instar (Profile 4, Fig. 5C). 212 genes were assigned to profile 10, which showed oscillating expression over time (Fig. 5A). GO-term enrichment in Profile 4 genes were associated with signal transduction and protein ubiquitination, but see figure 5D for a complete list. Genes assigned to each GO-term are reported in Table S3.

**Figure 5:**
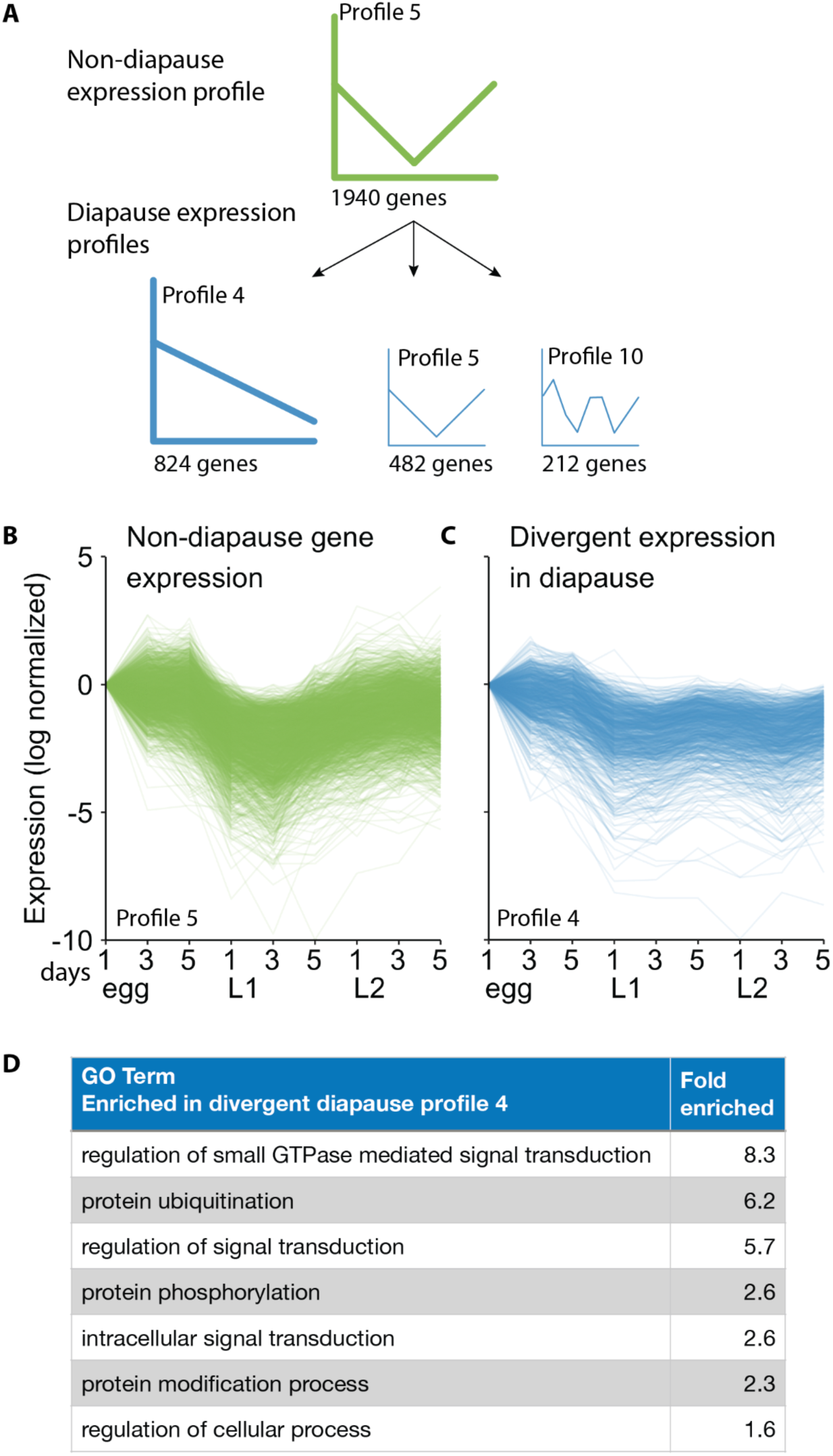
Divergence between the non-diapause temporal gene expression profile 5 and the diapause strain. A. Genes assigned to temporal expression profile 5 in the non-diapause strain were assigned to different profiles in the diapause strain. B. Non-diapause profile 5. C. Divergent genes assigned to diapause expression profile 4. D: GO term enrichment in the divergent gene expression profile. Fold enriched is the total number of genes over the expected number of genes assigned to a GO term.

In the third non-diapause profile (Profile 9), 674 genes exhibited expression that increased in expression in egg and early 1st instar larvae, but decreased in expression halfway through the first instar (Fig. 6A,B). Only 42 genes were assigned to the same profile in the diapausing strain, while 380 were assigned to profile 16 (continuous increase in expression over time) (Fig. 6C). GO-term enrichment included the glycolytic process, cellular catabolic processes and cellular signal transduction (see Fig. 6D for a complete list). Genes assigned to each GO-term are reported in Table S4.

**Figure 6:**
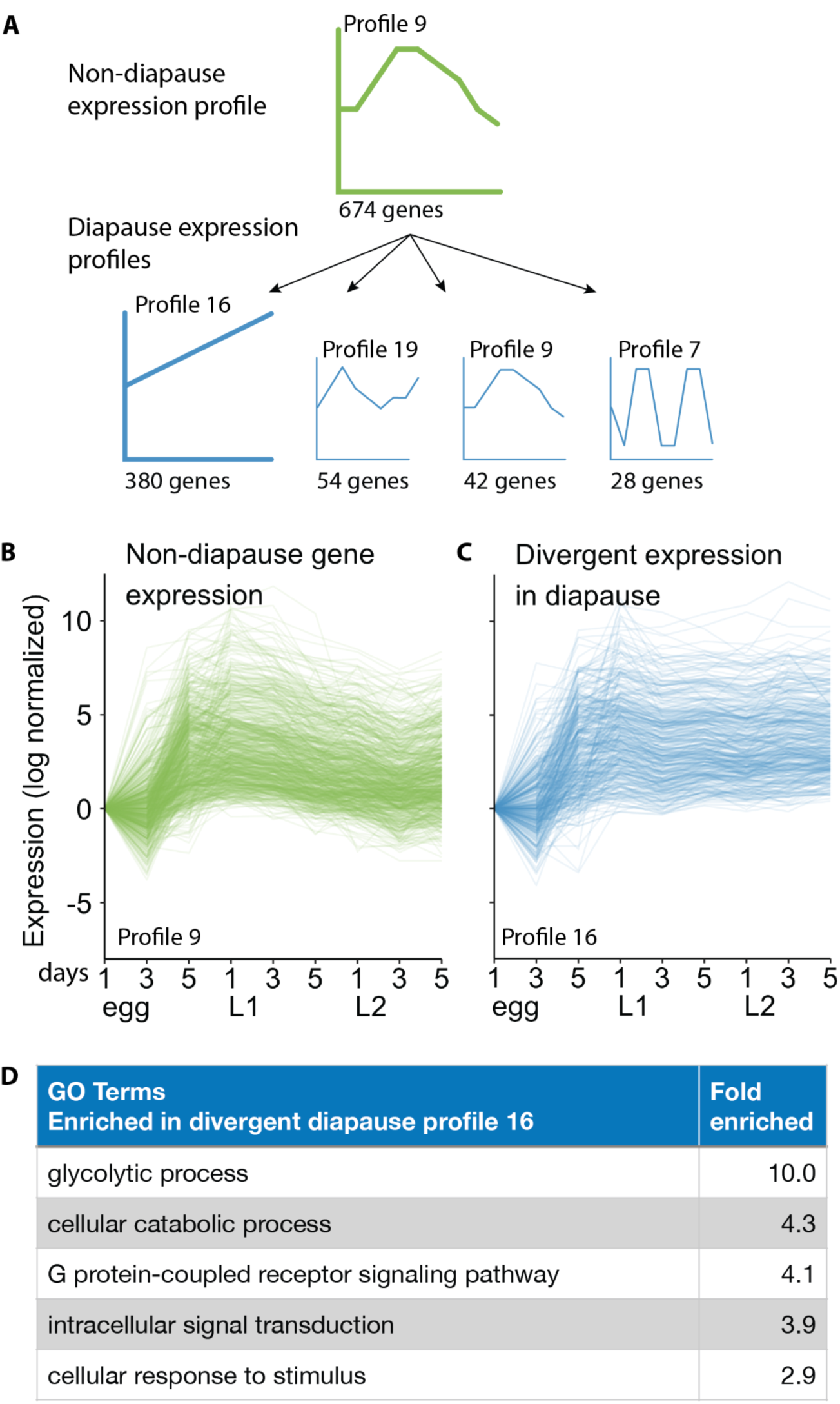
Divergence between the non-diapause temporal gene expression profile 9 and the diapause strain. A. Genes assigned to temporal expression profile 9 in the non-diapause strain were assigned to different profiles in the diapause strain. B. Non-diapause profile 9. C. Divergent genes assigned to diapause expression profile 16. D: GO term enrichment in the divergent gene expression profile. Fold enriched is the total number of genes over the expected number of genes assigned to a GO term.

Because the GO-terms for signal transduction (profile 4, decrease in diapause) and intracellular signal transduction (profile 16, increase in diapause) were very similar, and belonged to the same GO-term ancestor tree (SI Fig S2), we decided to further investigate the underlying genes for these terms. The genes associated with signal transduction that decreased in expression in the diapause strain were largely associated with growth and GTPase activity, whereas 40% of all genes associated with signal transduction that increased in expression in the diapause strain were involved with sensory signaling transduction (Table S5)

## Discussion

Diapause is a complex trait that requires extensive physiological changes in development, physiology and metabolism. Here, we characterize the genetic mechanisms regulating the non-diapause phenotype in a selected strain of spruce budworm, created through artificial selection on within-population variation of diapause incidence. We provide a chromosome-level genome assembly of the non-diapause strain, allowing direct comparison with a recently published genome of a diapause strain that was used to create the non-diapause strain. Genome comparison of the diapause and the non-diapause strains did not reveal evidence of major chromosomal rearrangements, nor did we see any evidence of a locus of major divergence between the strains. We did, however, reveal extensive transcriptional divergence starting approximately halfway through the first instar when we performed gene expression analysis of a fine-scale time-series comparing the non-diapause and diapause strains.

### Genetic divergence

We did not find any evidence of a single locus of large effect that could be responsible for causing a phenotypic shift from diapause to non-diapause development in these two strains. Were the shift in diapause incidence due to a chromosomal inversion, we would have found evidence of this with our whole genome alignments in the form of a break in the synteny between two chromosomes (Armstrong et al., 2019). We did not observe such breaks, and slight divergence from the diagonal is likely due to transposable elements. Further investigation of genomic divergence between the genomes included a sliding window analysis to identify SNP piles-ups, which would appear as an increase in nucleotide variants above the background signal. Artificial selection on a single locus of major effect is expected to result in a large number of nucleotide variants in one locus of the genome. When we tested for nucleotide divergence between the genomes, we did observe regions of high divergence, but all of these regions had high levels of repetitive sequences or had missing nucleotides nearby; both of which can result in an apparent localized accumulation of SNP variants. Although we cannot rule out that such regions are associated with the diapause phenotype, the elevated divergence in these regions is more likely to be an artifact than a real signal.

The lack of evidence for a single locus of large effect leads us to speculate that the change in diapause in our colonies is polygenic in nature. This would be in direct contrast with numerous other studies pointing towards one or a few genes associated with photoperiod signaling or other external signals (Cogni et al., 2014; Williams et al., 2006). However, these results identifying a single locus of large effect could be due to the nature of the analysis: QTL mapping tends to find only a few loci of large effect (Ragland et al., 2019). Indeed, when a whole genome association study was used to identify genetic variation associated with diapause in the speckled wood butterfly, *Pararge aegeria*, multiple loci of small effect were found in addition to two loci of large effect, indicating the polygenic nature of diapause phenotype (Pruisscher et al., 2018). To resolve the genetic architecture of the shift from diapause to non-diapause in the spruce budworm, a genome wide association study including many individuals will likely be the best approach.

### Transcriptomic divergence

We investigated gene expression at nine time points comparing diapause and non-diapause spruce budworm larvae from egg to immediately prior to diapause initiation. For both strains we saw a decrease of expression of genes associated with DNA-templated transcription, and a decrease followed by an increase in expression of genes associated with post translational modification (Fig 3). This change in gene expression activity is likely due to the general development from embryo to second instar larvae. The GO terms differed for profile 16 – increase in gene expression over time – where ‘glycerol-3-phosphate metabolic process’ was enriched in the diapause strain, and ‘lipid catabolic process’ was enriched in the non-diapause strain. This difference is very likely due to the differences in metabolism between the two strains, where diapausing spruce budworm do not feed, and rely entirely on glycolytic metabolism during overwintering (Han and Bauce, 1995; Marshall and Roe, 2021). In addition, diapausing spruce budworm start to produce a large amount of glycerol to increase cold-hardiness (Han and Bauce, 1995) (Fig. 1). This contrasts with the non-diapausing budworm that are starting to feed, and thus are upregulating genes associated with regular developmental metabolism. The other clear difference between the strains showed up for profile 19, with an increase-decrease-increase expression profile. In the diapause strain, this profile was associated with DNA-binding transcription activity, and the non-diapause strain profile was associated with the transmembrane receptor protein tyrosine kinase signaling pathway. There is no clear-cut hypothesis as to why these profiles are enriched for different GO terms, but it is most likely due to the difference in developmental programs between the strains.

Our gene expression results revealed extensive divergence between expression profiles; a large portion of genes assigned to one profile in the non-diapause strain were assigned to different profiles in the diapause strain. Genes that were upregulated in the diapause strain, but not in the non-diapause strain, showed a variety of enriched GO terms. For example, we found a strong enrichment of ‘negative regulation of macromolecule metabolic process,’ which is unsurprising given that downregulation of metabolism is common in diapausing insects. We also saw an upregulation of genes associated with the cellular response to stress, all of which also had the GO-term ‘DNA damage response’. This is very likely part of the preparation for diapause, where cold temperatures may lead to an increase in DNA damage and cellular stress (Somero, 2020). Another expected result was the divergent upregulation of expression of the glycolytic process. As mentioned, diapausing spruce budworm will produce a large amount of glycerol from stored glycogen to increase freeze resistance. Intriguingly, SNP variation in glycerol-3-phosphate dehydrogenase [NAD(+)] was identified in a linkage block that defined genomic differences among eastern and central eastern spruce budworm populations (Lumley et al. 2020). Given the role that this enzyme plays in glycerol metabolism (Storey and Storey, 2012), variation in the induction of cold tolerance invoked by the diapause program may exist among spruce budworm populations.

We were surprised to find a divergent increase in expression of genes associated with DNA replication and transcription in the diapause strain (Fig 4F). Usually, diapause is described as a state of transcriptional shutdown. A meta-analysis of 11 different transcriptomic studies comparing diapausing and non-diapausing insects found consistent downregulation of genes associated with transcription and cell cycle (Ragland and Keep, 2017), whereas we found an upregulation of these genes (eg. GO terms ‘DNA replication’ and ‘transcription by RNA polymerase II’). It is possible this discrepancy is due to the difference in sampling times, as our time series encompasses the pre-diapause and early diapause phase, and the meta-analysis focuses on insects already in diapause. A study in the mosquito *Aedes albopictus* (Poelchau et al., 2013) also showed an increase in expression of cell cycle associated genes in the early pre-diapause stage. It could be that we are seeing something similar, that in the early pre-diapause phase, cell cycling and transcription is actually increased to prepare for the oncoming period of arrested development.

Finally, we observed an upregulation of genes in diapausing spruce budworm associated with environmental cue processing, and more specifically, photoreception. Anecdotally, we have observed that increased light intensity affects diapause initiation in our spruce budworm colonies (Roe, unpublished). Interestingly, genetic variation in an ultraviolet-sensitive opsin was also included within the linkage block separating the eastern and central populations of the eastern spruce budworm (Lumley et al. 2020). These results could be related to the fact that photoperiod is a very common diapause signal. Gene expression and genetic mapping studies regularly point towards circadian clock genes as major factors in diapause evolution (Kozak et al., 2019; Pruisscher et al., 2018; Sandrelli et al., 2007; Tauber et al., 2007). Several other studies show a tight link between opsin expression and circadian clock genes (de Assis et al., 2016; Komada et al., 2015; Yan et al., 2014). In our study we did not detect an enrichment of circadian clock genes in our divergent expression analysis. However, given the tight link between opsins and circadian clock genes, there is a possible involvement of the circadian rhythm along with other light sensing mechanisms to alter diapause induction.

Surprisingly, the divergence between diapause and non-diapause gene expression profiles did not occur until halfway through the first instar, which suggests that the signal to induce the diapause preparation phase happens early in the first instar. This is reflected in the behavioral phenotype expressed during this period of development, where first instar larvae destined for diapause seek an overwintering site, spin a hibernaculum and clear their gut contents in preparation for diapause and the associated stress of winter (Han and Bauce, 1998). In most species, diapause is a responsive trait, where the signal to induce diapause comes from external conditions such as day length or temperature. During this sensitive period, the external signal is perceived and translated into an internal signal that initiates the diapause program (Kostál, 2006). In the case of the eastern spruce budworm, the second instar diapause is usually obligatory, where larvae will enter diapause regardless of external conditions. However, some variation exists in wild populations, where a small number of larvae will not enter diapause (Harvey, 1957). Also, the incidence of the non-diapause phenotype can be increased with early exposure to 24 hour light conditions (Cusson, Roe, unpublished), which indicates that diapause induction is responsive for at least part of the natural population. Taken together, there most likely is one type of signal that induces diapause, and it can rapidly evolve from ever-present, to responsive, to absent altogether. Future investigation into the molecular mechanisms of diapausing and non-diapausing strains of spruce budworm will help us understand the minimal changes necessary to alter the diapause phenotype in this widespread forest species.

## Acknowledgements

KEM is supported by an NSERC Discovery Grant. KB is supported by the NSF award DBI-2208932. Support was provided to ADR by funding from the Genomics Research and Development Initiative (GRDI-Government of Canada).

## Data Availability

The new genome assembly and all new data described in this article are available at NCBI under BioProject accession number PRJNA1173661.

## Notes

### Competing Interest Statement

The authors have declared no competing interest.

## Cited literature

Alonge, M., Lebeigle, L., Kirsche, M., Jenike, K., Ou, S., Aganezov, S., Wang, X., Lippman, Z.B., Schatz, M.C., Soyk, S., 2022. Automated assembly scaffolding using RagTag elevates a new tomato system for high-throughput genome editing. Genome Biol. 23, 258. 10.1186/s13059-022-02823-7

Armstrong, J., Fiddes, I.T., Diekhans, M., Paten, B., 2019. Whole-Genome Alignment and Comparative Annotation. Annu. Rev. Anim. Biosci. 7, 41–64. 10.1146/annurev-animal-020518-115005

Béliveau, C., Gagné, P., Picq, S., Vernygora, O., Keeling, C.I., Pinkney, K., Doucet, D., Wen, F., Spencer Johnston, J., Maaroufi, H., Boyle, B., Laroche, J., Dewar, K., Juretic, N., Blackburn, G., Nisole, A., Brunet, B., Brandão, M., Lumley, L., Duan, J., Quan, G., Lucarotti, C.J., Roe, A.D., Sperling, F.A.H., Levesque, R.C., Cusson, M., 2022. The Spruce Budworm Genome: Reconstructing the Evolutionary History of Antifreeze Proteins. Genome Biol. Evol. 14, evac087. 10.1093/gbe/evac087

Blum, M., Chang, H.-Y., Chuguransky, S., Grego, T., Kandasaamy, S., Mitchell, A., Nuka, G., Paysan-Lafosse, T., Qureshi, M., Raj, S., Richardson, L., Salazar, G.A., Williams, L., Bork, P., Bridge, A., Gough, J., Haft, D.H., Letunic, I., Marchler-Bauer, A., Mi, H., Natale, D.A., Necci, M., Orengo, C.A., Pandurangan, A.P., Rivoire, C., Sigrist, C.J.A., Sillitoe, I., Thanki, N., Thomas, P.D., Tosatto, S.C.E., Wu, C.H., Bateman, A., Finn, R.D., 2021. The InterPro protein families and domains database: 20 years on. Nucleic Acids Res. 49, D344. 10.1093/nar/gkaa977

Bolger, A.M., Lohse, M., Usadel, B., 2014. Trimmomatic: a flexible trimmer for Illumina sequence data. Bioinformatics 30, 2114. 10.1093/bioinformatics/btu170

Cogni, R., Kuczynski, C., Koury, S., Lavington, E., Behrman, E.L., O’Brien, K.R., Schmidt, P.S., Eanes, W.F., 2014. The intensity of selection acting on the couch potato gene--spatial-temporal variation in a diapause cline. Evol. Int. J. Org. Evol. 68, 538–548. 10.1111/evo.12291

Coombe, L., Li, J.X., Lo, T., Wong, J., Nikolic, V., Warren, R.L., Birol, I., 2021. LongStitch: high-quality genome assembly correction and scaffolding using long reads. BMC Bioinformatics 22, 534. 10.1186/s12859-021-04451-7

Dalla Benetta, E., Beukeboom, L.W., van de Zande, L., 2019. Adaptive Differences in Circadian Clock Gene Expression Patterns and Photoperiodic Diapause Induction in Nasonia vitripennis. Am. Nat. 193, 881–896. 10.1086/703159

de Assis, L.V.M., Moraes, M.N., da Silveira Cruz-Machado, S., Castrucci, A.M.L., 2016. The effect of white light on normal and malignant murine melanocytes: A link between opsins, clock genes, and melanogenesis. Biochim. Biophys. Acta BBA - Mol. Cell Res. 1863, 1119–1133. 10.1016/j.bbamcr.2016.03.001

De Coster, W., D’Hert, S., Schultz, D.T., Cruts, M., Van Broeckhoven, C., 2018. NanoPack: visualizing and processing long-read sequencing data. Bioinformatics 34, 2666–2669. 10.1093/bioinformatics/bty149

Denlinger, D.L., 2022. Insect Diapause. Cambridge University Press, Cambridge. 10.1017/9781108609364

Denlinger, D.L., 2002. Regulation of diapause. Annu Rev Entomol 47, 93–122. 10.1146/annurev.ento.47.091201.145137

Ebling, P.M., Dedes, J., 2015. IPS/003/003-Rearing diapause Choristoneura fumiferana. Standard Operating Procedure IPS/003/003.

Ernst, J., Bar-Joseph, Z., 2006. STEM: a tool for the analysis of short time series gene expression data. BMC Bioinformatics 7, 191. 10.1186/1471-2105-7-191

Fabian, D.K., Kapun, M., Nolte, V., Kofler, R., Schmidt, P.S., Schlötterer, C., Flatt, T., 2012. Genome-wide patterns of latitudinal differentiation among populations of *Drosophila melanogaster* from North America. Mol. Ecol. 21, 4748–4769. 10.1111/j.1365-294X.2012.05731.x

Ghurye, J., Rhie, A., Walenz, B.P., Schmitt, A., Selvaraj, S., Pop, M., Phillippy, A.M., Koren, S., 2019. Integrating Hi-C links with assembly graphs for chromosome-scale assembly. PLOS Comput. Biol. 15, e1007273. 10.1371/journal.pcbi.1007273

Gill, H.K., Goyal, G., Chahil, G., 2017. Insect Diapause: A Review. J. Agric. Sci. Technol. A 7. 10.17265/2161-6256/2017.07.002

Grabherr, M.G., Haas, B.J., Yassour, M., Levin, J.Z., Thompson, D.A., Amit, I., Adiconis, X., Fan, L., Raychowdhury, R., Zeng, Q., Chen, Z., Mauceli, E., Hacohen, N., Gnirke, A., Rhind, N., di Palma, F., Birren, B.W., Nusbaum, C., Lindblad-Toh, K., Friedman, N., Regev, A., 2011. Trinity: reconstructing a full-length transcriptome without a genome from RNA-Seq data. Nat. Biotechnol. 29, 644–652. 10.1038/nbt.1883

Grabherr, M.G., Russell, P., Meyer, M., Mauceli, E., Alföldi, J., Di Palma, F., Lindblad-Toh, K., 2010. Genome-wide synteny through highly sensitive sequence alignment: *Satsuma*. Bioinformatics 26, 1145–1151. 10.1093/bioinformatics/btq102

Han, E.-N., Bauce, E., 1998. Timing of diapause initiation, metabolic changes and overwintering survival of the spruce budworm, *Choristoneura fumiferana*. Ecol. Entomol. 23, 160–167. 10.1046/j.1365-2311.1998.00111.x

Han, E.-N., Bauce, E., 1995. Glycerol synthesis by diapausing larvae in response to the timing of low temperature exposure, and implications for overwintering survival of the spruce budworm, *Choristoneura fumiferana*. J. Insect Physiol. 41, 981–985. 10.1016/0022-1910(95)00049-Z

Harvey, G.T., 1961. Second Diapause in Spruce Budworm from Eastern Canada. Can. Entomol. 93, 594–602. 10.4039/Ent93594-7

Harvey, G.T., 1957. The occurrence and nature of diapause-free development in the spruce budworm, *Choristoneura fumiferana* (clem.) (lepidoptera: tortricidae). Can. J. Zool. 35, 549–572. 10.1139/z57-047

Holt, C., Yandell, M., 2011. MAKER2: an annotation pipeline and genome-database management tool for second-generation genome projects. BMC Bioinformatics 12, 491. 10.1186/1471-2105-12-491

Jones, P., Binns, D., Chang, H.-Y., Fraser, M., Li, W., McAnulla, C., McWilliam, H., Maslen, J., Mitchell, A., Nuka, G., Pesseat, S., Quinn, A.F., Sangrador-Vegas, A., Scheremetjew, M., Yong, S.-Y., Lopez, R., Hunter, S., 2014. InterProScan 5: genome-scale protein function classification. Bioinformatics 30, 1236–1240. 10.1093/bioinformatics/btu031

Komada, S., Kamae, Y., Koyanagi, M., Tatewaki, K., Hassaneen, E., Saifullah, A., Yoshii, T., Terakita, A., Tomioka, K., 2015. Green-sensitive opsin is the photoreceptor for photic entrainment of an insect circadian clock. Zool. Lett. 1, 11. 10.1186/s40851-015-0011-6

Koren, S., Walenz, B.P., Berlin, K., Miller, J.R., Bergman, N.H., Phillippy, A.M., 2017. Canu: scalable and accurate long-read assembly via adaptive k-mer weighting and repeat separation. Genome Res. 27, 722. 10.1101/gr.215087.116

Korf, I., 2004. Gene finding in novel genomes. BMC Bioinformatics 5, 59. 10.1186/1471-2105-5-59

Kostál, V., 2006. Eco-physiological phases of insect diapause. J. Insect Physiol. 52, 113–127. 10.1016/j.jinsphys.2005.09.008

Kozak, G.M., Wadsworth, C.B., Kahne, S.C., Bogdanowicz, S.M., Harrison, R.G., Coates, B.S., Dopman, E.B., 2019. Genomic Basis of Circannual Rhythm in the European Corn Borer Moth. Curr. Biol. 29, 3501–3509.e5. 10.1016/j.cub.2019.08.053

Kurtz, S., Phillippy, A., Delcher, A.L., Smoot, M., Shumway, M., Antonescu, C., Salzberg, S.L., 2004. Versatile and open software for comparing large genomes. Genome Biol.

Liao, Y., Smyth, G.K., Shi, W., 2019. The R package Rsubread is easier, faster, cheaper and better for alignment and quantification of RNA sequencing reads. Nucleic Acids Res. 47, e47. 10.1093/nar/gkz114

Manni, M., Berkeley, M.R., Seppey, M., Zdobnov, E.M., 2021. BUSCO: Assessing Genomic Data Quality and Beyond. Curr. Protoc. 1, e323. 10.1002/cpz1.323

Marshall, K.E., Roe, A.D., 2021. Surviving in a Frozen Forest: the Physiology of Eastern Spruce Budworm Overwintering. Physiology 36, 174–182. 10.1152/physiol.00037.2020

Martin, M., 2011. Cutadapt removes adapter sequences from high-throughput sequencing reads. EMBnet.journal 17, 10–12. 10.14806/ej.17.1.200

Mathias, D., Jacky, L., Bradshaw, W.E., Holzapfel, C.M., 2007. Quantitative Trait Loci Associated with Photoperiodic Response and Stage of Diapause in the Pitcher-Plant Mosquito, *Wyeomyia smithii*. Genetics 176, 391–402. 10.1534/genetics.106.068726

McMorran, A., 1965. A Synthetic Diet for the Spruce Budworm, *Choristoneura fumiferana* (Clem.) (Lepidoptera: Tortricidae). Can. Entomol. 97, 58–62. 10.4039/Ent9758-1

Omni-C 0.1 documentation [WWW Document], n.d. URL https://omni-c.readthedocs.io (accessed 10.17.24).

Poelchau, M.F., Reynolds, J.A., Elsik, C.G., Denlinger, D.L., Armbruster, P.A., 2013. Deep sequencing reveals complex mechanisms of diapause preparation in the invasive mosquito, Aedes albopictus. Proc. R. Soc. B Biol. Sci. 280, 20130143. 10.1098/rspb.2013.0143

Pruisscher, P., Nylin, S., Gotthard, K., Wheat, C.W., 2018. Genetic variation underlying local adaptation of diapause induction along a cline in a butterfly. Mol. Ecol. 27, 3613–3626. 10.1111/mec.14829

Ragland, G.J., Armbruster, P.A., Meuti, M.E., 2019. Evolutionary and functional genetics of insect diapause: a call for greater integration. Curr. Opin. Insect Sci., Neuroscience • Special section on Evolutionary Genetics and Genomics 36, 74–81. 10.1016/j.cois.2019.08.003

Ragland, G.J., Denlinger, D.L., Hahn, D.A., 2010. Mechanisms of suspended animation are revealed by transcript profiling of diapause in the flesh fly. Proc. Natl. Acad. Sci. 107, 14909–14914. 10.1073/pnas.1007075107

Ragland, G.J., Keep, E., 2017. Comparative transcriptomics support evolutionary convergence of diapause responses across Insecta. Physiol. Entomol. 42, 246–256. 10.1111/phen.12193

Régnière, J., Duval, P., 1998. Overwintering mortality of spruce budworm, *Choristoneura fumiferana* (Clem.) (Lepidoptera: Tortricidae) populations under field conditions. Can. Entomol. 130, 13–26. 10.4039/Ent13013-1

Rio, D.C., Ares, M., Hannon, G.J., Nilsen, T.W., 2010. Purification of RNA using TRIzol (TRI reagent). Cold Spring Harb. Protoc. 2010, pdb.prot5439. 10.1101/pdb.prot5439

Roe, A.D., Demidovich, M., Dedes, J., 2018. Origins and History of Laboratory Insect Stocks in a Multispecies Insect Production Facility, With the Proposal of Standardized Nomenclature and Designation of Formal Standard Names. J. Insect Sci. 18, 1. 10.1093/jisesa/iey037

Roe, A.D., Wardlaw, A.A., Butterson, S., Marshall, K.E., 2024. Diapause survival requires a temperature-sensitive preparatory period. Curr. Res. Insect Sci. 5, 100073. 10.1016/j.cris.2024.100073

Sandrelli, F., Tauber, E., Pegoraro, M., Mazzotta, G., Cisotto, P., Landskron, J., Stanewsky, R., Piccin, A., Rosato, E., Zordan, M., Costa, R., Kyriacou, C.P., 2007. A Molecular Basis for Natural Selection at the timeless Locus in *Drosophila melanogaster*. Science 316, 1898– 1900. 10.1126/science.1138426

Somero, G.N., 2020. The cellular stress response and temperature: Function, regulation, and evolution. J. Exp. Zool. Part Ecol. Integr. Physiol. 333, 379–397. 10.1002/jez.2344

Stanke, M., Morgenstern, B., 2005. AUGUSTUS: a web server for gene prediction in eukaryotes that allows user-defined constraints. Nucleic Acids Res. 33, W465–467. 10.1093/nar/gki458

Storey, K.B., Storey, J.M., 2012. Insect cold hardiness: metabolic, gene, and protein adaptation ^1^ This review is part of a virtual symposium on recent advances in understanding a variety of complex regulatory processes in insect physiology and endocrinology, including development, metabolism, cold hardiness, food intake and digestion, and diuresis, through the use of omics technologies in the postgenomic era. Can. J. Zool. 90, 456–475. 10.1139/z2012-011

Supek, F., Bošnjak, M., Škunca, N., Šmuc, T., 2011. REVIGO Summarizes and Visualizes Long Lists of Gene Ontology Terms. PLOS ONE 6, e21800. 10.1371/journal.pone.0021800

Tauber, E., Zordan, M., Sandrelli, F., Pegoraro, M., Osterwalder, N., Breda, C., Daga, A., Selmin, A., Monger, K., Benna, C., Rosato, E., Kyriacou, C.P., Costa, R., 2007. Natural selection favors a newly derived timeless allele in *Drosophila melanogaster*. Science 316, 1895–1898. 10.1126/science.1138412

The UniProt Consortium, 2023. UniProt: the Universal Protein Knowledgebase in 2023. Nucleic Acids Res. 51, D523–D531. 10.1093/nar/gkac1052

Tyukmaeva, V.I., Salminen, T.S., Kankare, M., Knott, K.E., Hoikkala, A., 2011. Adaptation to a seasonally varying environment: a strong latitudinal cline in reproductive diapause combined with high gene flow in Drosophila montana. Ecol. Evol. 1, 160–168. 10.1002/ece3.14

Walker, B.J., Abeel, T., Shea, T., Priest, M., Abouelliel, A., Sakthikumar, S., Cuomo, C.A., Zeng, Q., Wortman, J., Young, S.K., Earl, A.M., 2014. Pilon: An Integrated Tool for Comprehensive Microbial Variant Detection and Genome Assembly Improvement. PLoS ONE 9, e112963. 10.1371/journal.pone.0112963

Williams, K.D., Busto, M., Suster, M.L., So, A.K.-C., Ben-Shahar, Y., Leevers, S.J., Sokolowski, M.B., 2006. Natural variation in *Drosophila melanogaster* diapause due to the insulin-regulated PI3-kinase. Proc. Natl. Acad. Sci. U. S. A. 103, 15911–15915. 10.1073/pnas.0604592103

Wilsterman, K., Ballinger, M.A., Williams, C.M., 2021. A unifying, eco-physiological framework for animal dormancy. Funct. Ecol. 35, 11–31. 10.1111/1365-2435.13718

Wolff, J., Rabbani, L., Gilsbach, R., Richard, G., Manke, T., Backofen, R., Grüning, B.A., 2020. Galaxy HiCExplorer 3: a web server for reproducible Hi-C, capture Hi-C and single-cell Hi-C data analysis, quality control and visualization. Nucleic Acids Res. 48, W177– W184. 10.1093/nar/gkaa220

Yan, S., Zhu, J., Zhu, W., Zhang, X., Li, Z., Liu, X., Zhang, Q., 2014. The Expression of Three Opsin Genes from the Compound Eye of *Helicoverpa armigera* (Lepidoptera: Noctuidae) Is Regulated by a Circadian Clock, Light Conditions and Nutritional Status. PLOS ONE 9, e111683. 10.1371/journal.pone.0111683

