## Supplemental Figures 1-2 for "Major changes in gene expression between strains of diapausing and non-diapause spruce budworm despite little evidence of genetic divergence"

### SI figures

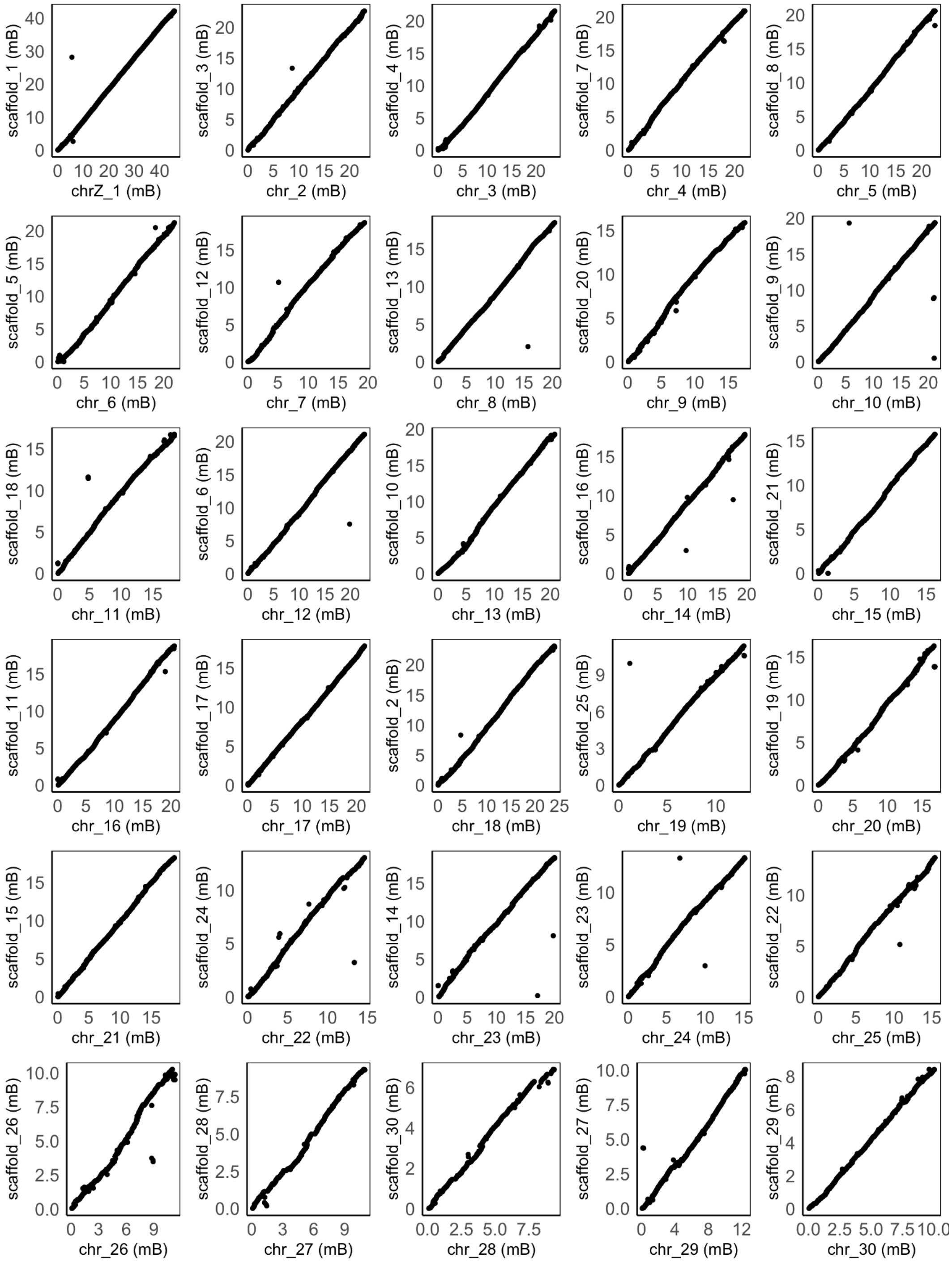


Supplemental figure 1.

**Chromosome alignment of the diapause (x-axis) and non-diapause (y-axis) genome.**

No major chromosome inversions were detected.

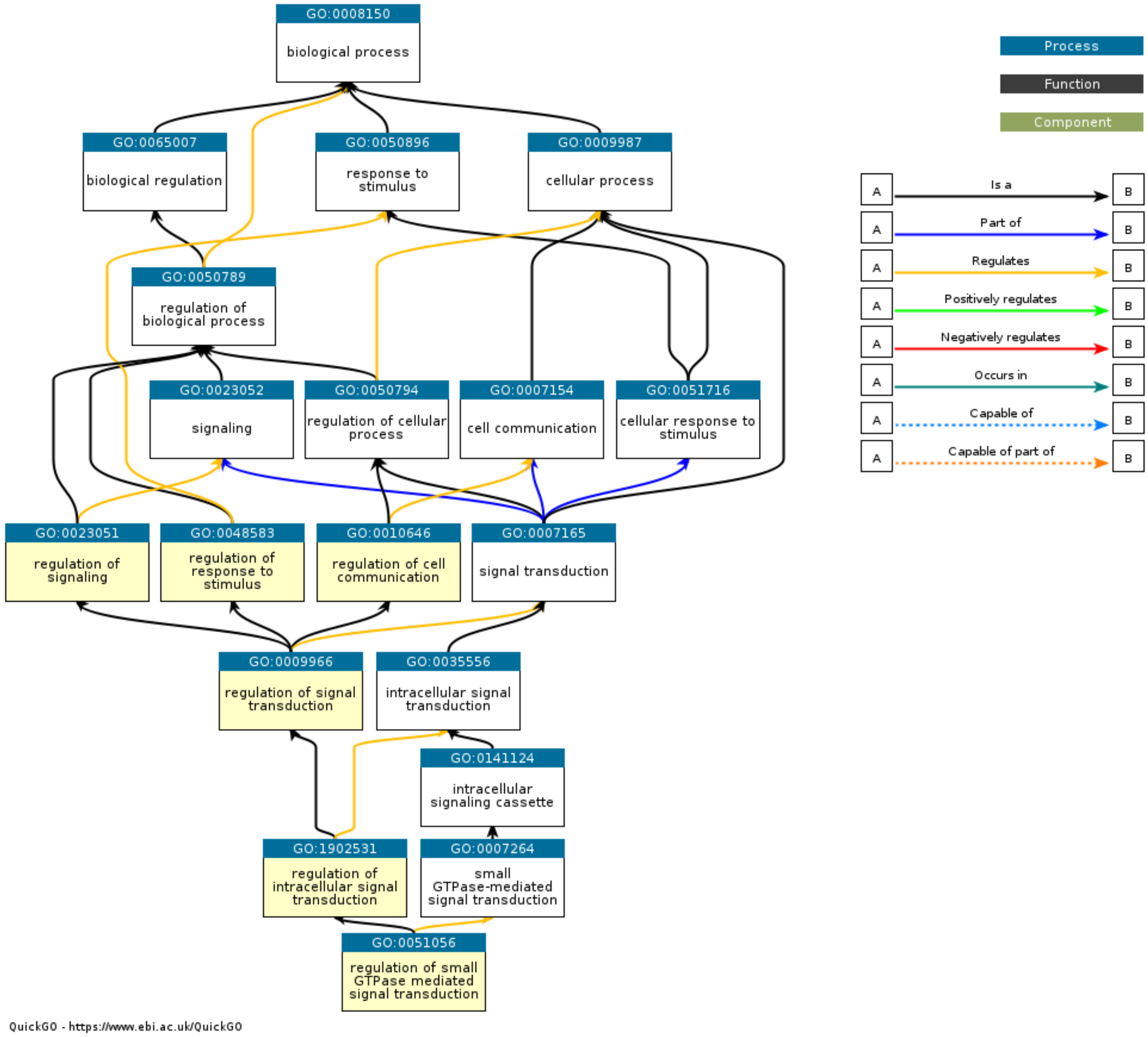


Supplemental figure 2

GO-terms for regulation of signal transduction (enriched in divergent profile 4, fig 5) and intracellular signal transduction (enriched in divergent profile 16, fig 4) were very similar, and belonged to the same GO-term ancestor tree.
